# The chromatin remodeler DEK promotes proliferation of mammary epithelium and is associated with H3K27me3 epigenetic modifications

**DOI:** 10.1101/2024.09.09.612116

**Authors:** Megan Johnstone, Ashley Leck, Taylor Lange, Katherine Wilcher, Miranda S. Shephard, Aditi Paranjpe, Sophia Schutte, Susanne Wells, Ferdinand Kappes, Nathan Salomonis, Lisa M. Privette Vinnedge

## Abstract

The DEK chromatin remodeling protein was previously shown to confer oncogenic phenotypes to human and mouse mammary epithelial cells using *in vitro* and knockout mouse models. However, its functional role in normal mammary gland epithelium remained unexplored. We developed two novel mouse models to study the role of Dek in normal mammary gland biology *in vivo*. Mammary gland-specific Dek over-expression in mice resulted in hyperproliferation of cells that visually resembled alveolar cells, and a transcriptional profile that indicated increased expression of cell cycle, mammary stem/progenitor, and lactation-associated genes. Conversely, Dek knockout mice exhibited an alveologenesis or lactation defect, resulting in dramatically reduced pup survival. Analysis of previously published single-cell RNA-sequencing of mouse mammary glands revealed that *Dek* is most highly expressed in mammary stem cells and alveolar progenitor cells, and to a lesser extent in basal epithelial cells, supporting the observed phenotypes. Mechanistically, we discovered that Dek is a modifier of Ezh2 methyltransferase activity, upregulating the levels of histone H3 trimethylation on lysine 27 (H3K27me3) to control gene transcription. Combined, this work indicates that Dek promotes proliferation of mammary epithelial cells via cell cycle deregulation. Furthermore, we report a novel function for Dek in alveologenesis and histone H3 K27 trimethylation.

## Introduction

Developmental pathways that maintain stem and progenitor cell pools, or promote cellular proliferation and migration during embryogenesis, are frequently re-activated and deregulated later in life during tumorigenesis.^[1]^ As an example, breast cancer is often characterized, in part, by the dysregulation of cell cycle (i.e., E2F and CDK4/6), DNA repair (i.e., BRCA1 and p53), and epigenetic modifying (i.e., EZH2) proteins.^[2–5]^ Additionally, chromatin remodeling and epigenetic modification pathways that are important for controlling cellular differentiation are also deregulated during tumorigenesis.^[6, 7]^

There is a substantial body of work on the role of the DEK protein in solid and hematological malignancies; however, its roles(s) in normal development are poorly understood. DEK is a chromatin remodeling protein and histone H3 chaperone.^[8–10]^ In *Drosophila*, transgenic models over-expressing Dek in the eye resulted in a rough eye phenotype due to caspase-mediated apoptosis. This correlated with hypoacetylation of lysine residues in histones H3 and H4, including H3K27, H3K9, and H4K5, resulting in epigenetic silencing of the anti-apoptotic gene *bcl-2*.^[11]^ In tissue-specific knockout mouse models, *Dek* loss impaired the self-renewing ability of hematopoietic stem cells by limiting quiescence and accelerating mitochondrial metabolism. This led to decreased bone marrow cellularity and loss of hematopoietic stem and progenitor cells that was associated with increased H3K27ac epigenetic marks. This resulted in the transcriptional deregulation of several genes related to proliferation and metabolism, including upregulated *Akt1/2, Ccnb1/2* (cyclin B)*, Cdkn1a* (p21), and *Cdkn1b* (p27). The role of Dek in directing histone acetylation was found to occur through the recruitment of co-repressor NCoR1 and inhibition of histone acetyltransferases p300 and PCAF.^[12]^ In addition, Dek deficient primary mouse neurons cultured *in vitro* demonstrated an increase in acetylation of lysine 36 of histone H3 (H3K36ac).^[13]^ These reports indicate that DEK suppresses histone H3 acetylation, leaving these residues available for methylation. Indeed, Dek was previously shown in *Drosophila* to support histone H3 trimethylation on residue lysine 9 (H3K9me3).^[14]^ However, mechanisms whereby DEK promotes histone H3 methylation, and biological processes wherein this is important, remain unexplored. Histone methylation, particularly on histone H3, is both dynamic and tightly regulated during development and is important for establishing cell lineages, body patterning, and the development of specific organs. Histone methylation is accomplished by a family of methyltransferases, such as EZH2 of the polycomb repressive complex 2 (PRC2), and can be reversed by histone demethylases.^[15]^

We, and others, have reported that DEK mRNA and protein are over-expressed in many solid cancers, including 62-92% of breast cancers.^[16–19]^ DEK has no known enzymatic functions but is a chromatin-modifying phosphoprotein with molecular activities in DNA repair, mRNA transcription, splicing, and histone modification.^[20–26]^ We previously reported *in vitro* studies using MCF10A immortalized human mammary epithelial cells where we demonstrated that DEK over-expression (DEK-OE) promotes cellular proliferation and invasion in 2D cultures and 3D organoid models.^[17, 27]^ In 3D organotypic models of skin, DEK-OE promoted hyperplasia as evidenced by epidermal thickening and increased expression of proliferation-associated markers like PCNA.^[28]^ A squamous epithelium-specific Dek transgenic mouse model indicated that Dek over-expression did not promote hyperplasia in control conditions but did increase esophageal tumor formation in a 4NQO chemical carcinogenesis model.^[29]^ However, this differed from a *Drosophila* transgenic model, where Dek-OE promoted apoptosis^[11]^ and in the mouse hematopoietic system, where Dek loss, not over-expression, promoted cell cycle entry and suppressed quiescence.^[12, 30]^ Therefore, the role of DEK in promoting, or inhibiting, cellular proliferation has not been consistent across model systems. Furthermore, the impact of deregulated DEK expression on normal mammary epithelium has not been previously reported.

We sought to define the transcriptional and functional consequences of Dek-OE in murine mammary epithelium. To this end, we generated a novel doxycycline-regulated mammary epithelium specific Dek-OE model, MMTV-tTA/BiLDek (Dek-OE) mice and analyzed transcriptomic consequences of Dek-OE in mammary glands by RNA-sequencing. We report that Dek over-expression is sufficient to cause mammary epithelial hyperplasia characterized by excessive proliferation and cell cycle deregulation and promotes expression of proteins associated with the luminal alveolar cell identity and milk production. We also report a novel Dek conditional knockout (Dek-cKO) mouse model with evidence of a primary lactation deficiency phenotype. Importantly, we report for the first time that Dek promotes H3K27me3 histone trimethylation in mammary epithelial cells *in vivo* and *in vitro* and physically interacts with EZH2 and other members of the PRC2 complex.

## Results

### Generation of mammary gland specific Dek over-expression in mice

To study the effects of Dek-OE on mammary epithelium, we created a novel, tetracycline (doxycycline) responsive transgenic model. Mouse Mammary Tumor Virus-tetracycline transactivator (MMTV-tTA) mice^[31]^ were crossed with the Bi-L-Dek mouse^[29]^ resulting in a MMTV-tTA/Bi-L-DEK (DEK-OE) mouse (Fig. 1A.) In this model, doxycycline repressed the transcription of the *Dek* transgene and the luciferase reporter via a bidirectional promotor containing the tetracycline response element (TRE). Representative genotyping for the *tTA*, luciferase, and *Dek* transgenes and the endogenous *Dek* gene is shown in Fig 1B. Expression of the luciferase reporter transgene with repression by doxycycline chow (DOX) was validated by *in vivo* imaging system (IVIS, Fig 1C). We detected approximately 2-fold higher Dek protein expression in whole mammary gland protein lysates and via immunohistochemical staining in tissues collected from Dek-OE virgin females in diestrus compared to Dox repressed (“control”) mice (Fig 1D-G). This is similar to the degree of OE observed in other *in vitro* and *in vivo* models.^[29, 32]^ Since mammary tumorigenesis can be a long process and is observed primarily in women over 50 years of age,^[33]^ we aged multiparous, non-lactating females to the human equivalent age, which is 15 months,^[34]^ to detect the impact of prolonged Dek-OE on mammary epithelium. Spontaneous tumor development was not observed during the 15 months of aging. However, we noted substantial hyperplasia in inguinal mammary glands marked by increased epithelial density, as quantified by Sholl analysis (Fig 1H-I).

**Fig 1.**
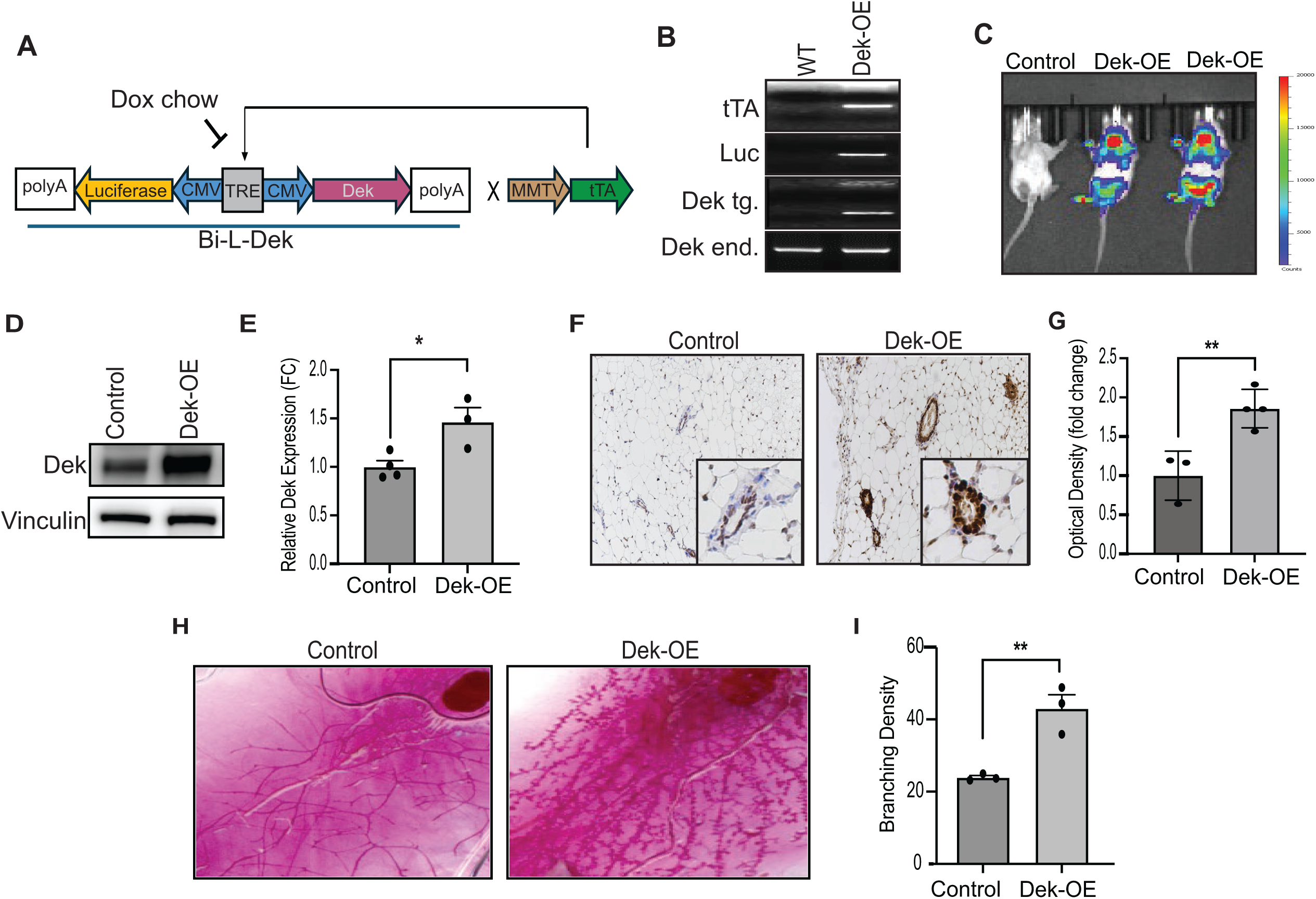
Generation of a tetracycline-off regulated mammary gland specific overexpression of *Dek* transgenic murine model. (A)Transgenic construct of Bi-L-Dek transgenic mice with a tetracycline response element (TRE) drives dual cytomegalovirus (CMV) promotors that bi-directionally transcribe luciferase and Dek. This transgene is transcribed by a tetracycline transactivator under the control of a murine mammary tumor virus (MMTV) promotor (MMTV-tTA), resulting in targeted mammary epithelium specific transgenic Dek over expression (Dek-OE). (B) Representative genotyping reveals presence of transgenic constructs described in A and presence of endogenous (WT) Dek (C) Examination of luciferase expression and tetracycline (doxycycline) repression in Dek-OE mice confirmed via after intraperitoneal injection of luciferin and visulized with an *in vivo* imaging system (IVIS). (D) Western blot analysis of Dek protein from whole lysed mammary gland. Vinculin served as a control for protein loading. (E) Western blot densitometry analysis reveals a 1.5-2-fold increase in DEK protein expression (n=4,3, p < 0.05). (F) Dek expression is elevated in Dek-OE mammary glands compared to +dox controls, as detected by immunohistochemistry. (G) Quantification of Dek immunohistochemistry from (F). Image J color deconvolution was used to measure staining intensity and is graphed as a fold-change (n=3,4, p<0.05). (H) Representative whole mount images from +dox control and Dek-OE inguinal mammary glands. (I) Image J was used to perform Sholl analysis and quantify epithelial branching density of glands from 60-week-old virgin females. (n=3,3 p<0.05). An unpaired Student’s t-test was used for statistical analysis and data is presented as mean +/− SEM.

### RNA sequencing reveals differential expression of cell cycle related genes

To discover molecular targets of Dek-OE we performed RNA sequencing on whole mammary tissue from two +dox control and two Dek-OE adult multiparous, non-lactating females at 15 months of age, when hyperplasia was observed. Principle component analysis (PCA) showed that the Dek-OE transcriptome is unique and distinct from control samples (Fig 2A). We plotted differentially expressed genes (DEGs) by volcano plot and found that 1631 genes were up-regulated and only 340 genes were down-regulated (Fig 2B). *Dek* mRNA over-expression in samples used for RNA-Seq was confirmed by plotting the FPKM for all samples (Fig 2C).

**Fig 2.**
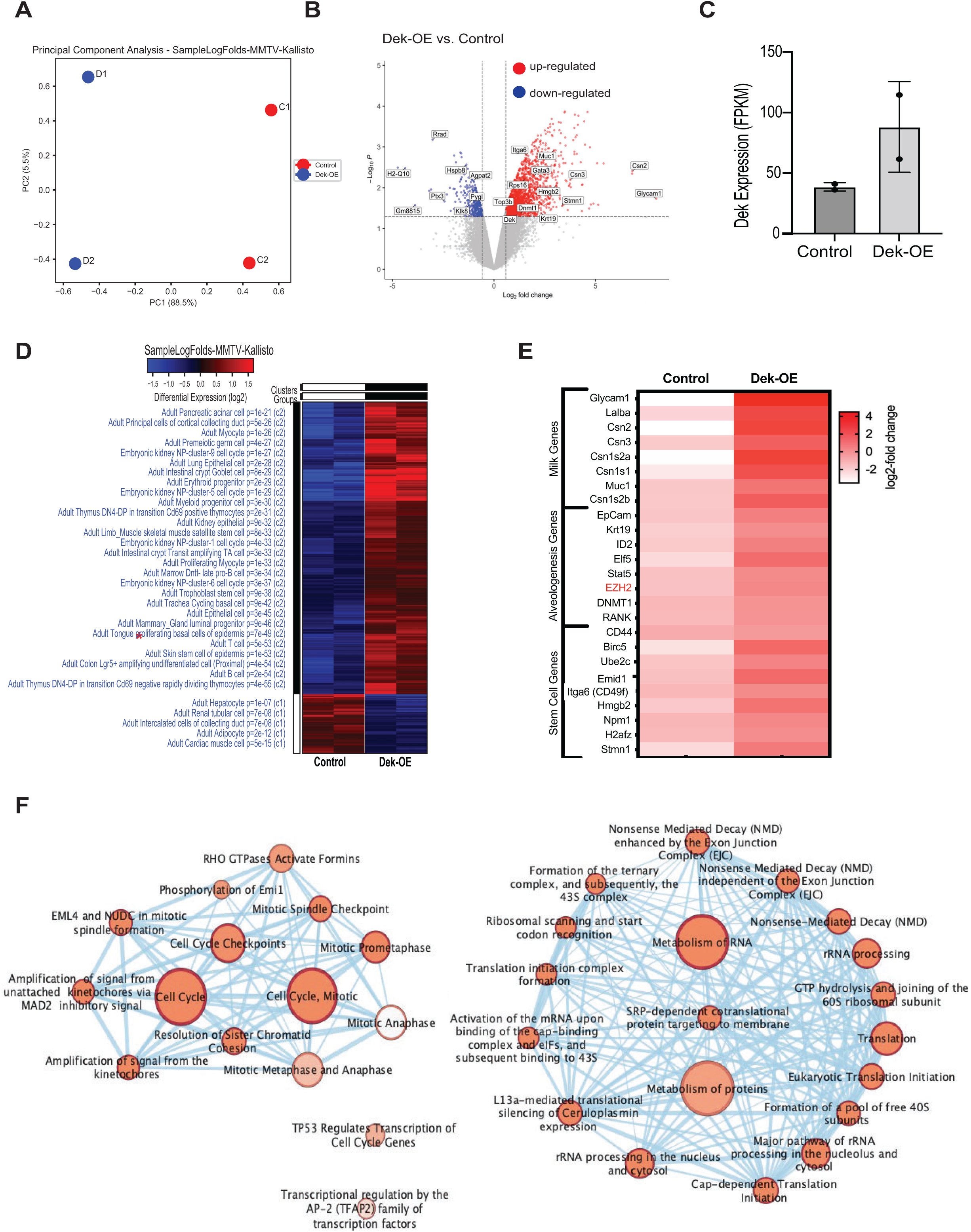
RNA sequencing reveals differential expression of cell cycle regulatory signaling and markers of mammary luminal progenitor cells. (A.) Principal component analysis (PCA) reveals that Dek-OE samples are distinct and unique from control samples (B) Volcano plot is used to visualize significant differentially up- and down regulated gene targets. Upregulated targets are depicted in red and down-regulated targets in blue. (C) *Dek* mRNA levels of samples used for bulk RNA-Seq (D) A heat map depicts the up- and down-regulated genes in Dek-OE mammary glands compared to +dox chow controls. Top hits of the most significant gene ontologies associated with the up- and down-regulated genes are listed on the left. * Note the ontology of “adult mammary gland luminal progenitor” p=9e^−46^. (E) A visual representation of a select set of genes strongly over-expressed in Dek-OE mammary glands compared to +dox controls that are indicative of an increase in luminal alveolar progenitor cells. Genes associated with milk protein production, alveologenesis, and stem cells are shown. (F) Significant categories of upregulated genes in Dek-OE mammary glands reveals a strong focus on cell cycle regulation, metabolism of RNA and proteins, and transcriptional regulation of AP-2 and p53 target genes. Nodes were identified using Cytoscape reactome analysis.

DEGs were plotted as a heatmap and ontologies for biomarkers of cell populations were defined. Up-regulated DEGs reflected cell biomarker signatures for several types of epithelial cells. Most notably, a specific ontology for “adult mammary gland luminal progenitor” was identified in Dek-OE samples (Fig 2D). Indeed, we observed an upregulation of genes associated with luminal alveolar progenitor mammary epithelial cells, including genes encoding milk proteins (Glycam1, Muc1, and caseins), luminal alveolar cell specific genes (Elf5, EpCam, Krt19, and EZH2), and stem/progenitor cell markers (CD44, Birc5/survivin, and Itga6/CD49f) (Fig 2B, 2E). We then used GO-Elite to identify transcription factors Yy1, Nfkb2, p53, and E2F as potentially responsible for the expression of Dek-OE-induced differentially expressed genes (Fig S1A) while transcription factors associated with the expression of down-regulated genes were limited, but included Pparg, Myod1, and Srebf1 (Fig S2A). We also used GO-Elite to assess KEGG pathways associated with DEGs caused by Dek over-expression. For upregulated genes, we identified p53 signaling, cell cycle, tight junctions and adherens junctions, ribosomes, and progesterone signaling (Fig S1B). For down-regulated genes, we identified KEGG pathways related to adipocytokine signaling and a small number of genes related to various metabolic pathways like glycolysis, citrate cycle, and pentose phosphate pathways (Fig S2B). Next, we used gProfiler to create a reactome network of upregulated genes in Dek-OE mammary glands, which indicated “cell cycle” and “metabolism of RNA/proteins” as central processes deregulated in response to Dek-OE (Fig 2F). The observed gene expression profile indicates a possible Dek dependent regulation of cellular processes relevant to mammary gland development, including cell cycle progression, metabolism, and progesterone signaling, that may promote hyperplasia in Dek-OE mammary epithelium.

One of the strongest gene ontology associations was “cell cycle processes” so we validated the respective DEGs with a focus on cyclins, cyclin-dependent kinases, and cell cycle inhibitors. Immununohistochemistry staining of 8-week old inguinal glands showed that expression levels of cell cycle inhibitors p21 and p27, which inhibit cyclin-CDK complexes, were significantly down-regulated with Dek-OE (Fig 3A-B). Cyclin A and Cdk proteins -2, -4, and -6 were also found to be highly expressed in Dek-OE mammary epithelium compared to controls (Fig 3C-3F). To test if Dek-mediated transcriptional deregulation contributed to hyperplasia, we isolated primary mammary epithelial cells from control and Dek-OE juvenile mice and cultured them as 3D organoids. Indeed, Dek-OE organoids were nearly three times larger in volume compared to controls. Importantly, *in vitro* hyperplasia was completely rescued by treating Dek-OE organoids with the CDK4/6 inhibitor palbociclib, an FDA-approved small molecule inhibitor used to treat estrogen receptor positive (ER+) breast cancer (Fig 3G-H). Palbociclib treatment did not reduce the volume of control cells, thus showing specificity for DEK-OE cells. The observed Dek-driven regulation of cell cycle effectors was sufficient to create a pro-proliferative phenotype. As early as 6 weeks of age, we detected an increased proportion of Ki67 positive proliferative cells in Dek-OE mammary epithelium (Fig 3I-J). Of note, the association between DEK and a pro-proliferative gene signature was not limited to mouse mammary epithelium. Using the METABRIC dataset in The Cancer Genome Atlas, we determined that *DEK* mRNA expression was inversely correlated with *CDKN1A* (p21) and positively correlated with *CCNA2* (cyclin A), *MKi67* (Ki67) and *PCNA* (Fig S3). This suggests that transcriptionally mediated consequences of Dek-OE in murine models may be relevant in humans.

**Fig 3.**
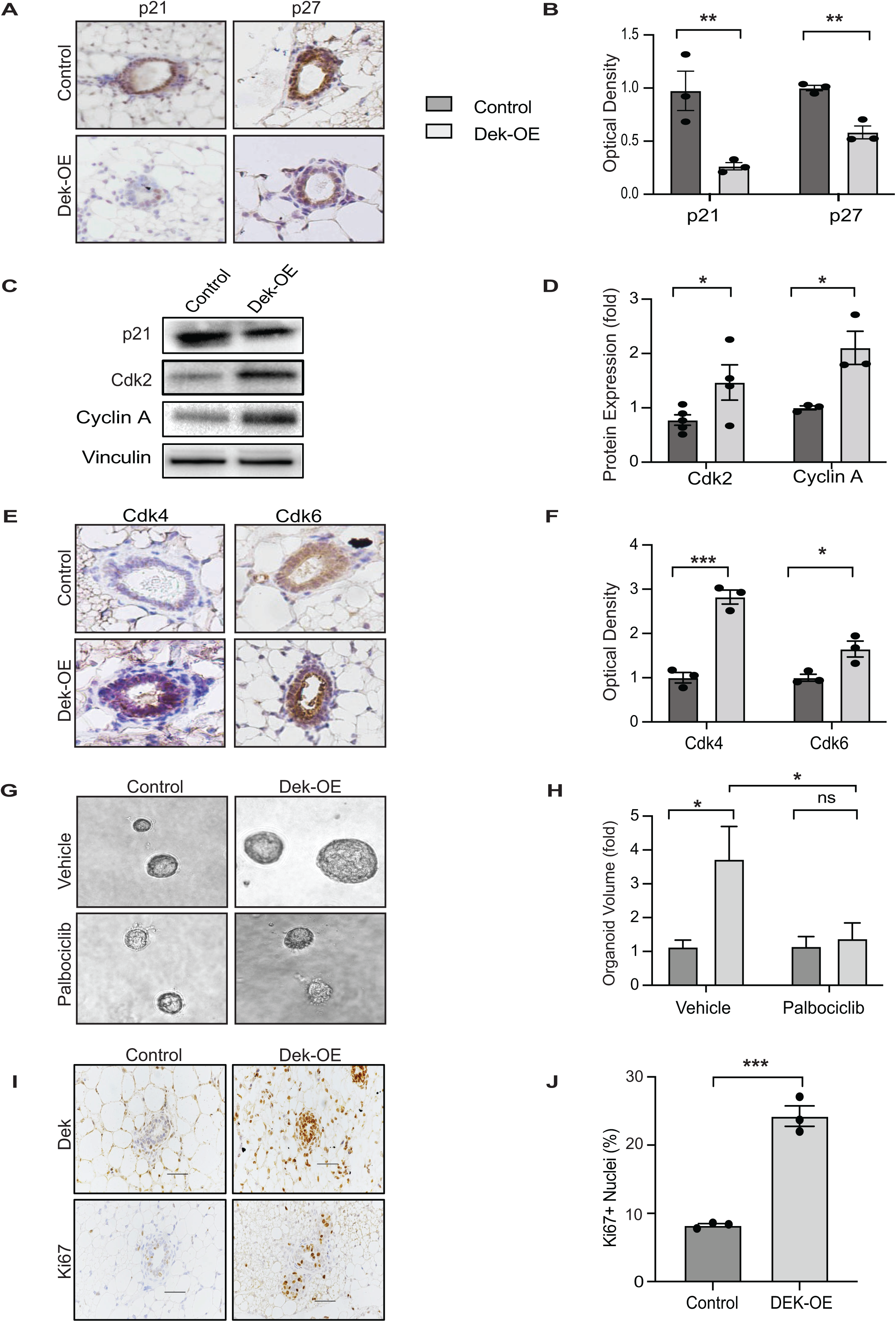
Dek-OE mice display a mitogenic phenotype associated with increased expression of cyclin-CDK complex proteins. (A) Dek-OE mammary epithelium has decreased expression of cell cycle inhibitor proteins p21 and p27 as detected by immunohistochemistry (B) Quantification of immunohistochemical staining in (A) using Image J color deconvolution and represented as fold change. (C) Representative western blot images showing decreased p21 protein expression and increased expression of Cdk2 and cyclin A in Dek-OE mammary glands compared to +dox controls. Vinculin served as protein loading controls (D) Quantification of Western blots in (C) using densitometry analysis. (E) Dek-OE mammary glands express more Cdk4 and Cdk6 compared to control glands as detected by immunohistochemistry. (F) Quantification of immunohistochemical staining in (E) using Image J color deconvolution to measure optical density, graphed as fold-change compared to +dox control samples. (G) Palbociclib, a Cdk4/6 small molecule inhibitor, prevents Dek-induced hyperplasia. Primary mammary epithelial cells were isolated and grown as 3D organoids with, or without, doxycycline or Palbociclib. Dek-OE organoids are 3-fold larger than +dox control samples, which is completely rescued with Palbociclib treatment. (H) Quantification of organoid volume using measurements obtained with Image J. (I) Dek-OE mammary glands have significantly more proliferating mammary epithelial cells as determined by Ki67 immunohistochemical staining. (J) Quantification of data presented in (I), with the percentage of Ki67+ nuclei within mammary epithelium shown. Sample size is ≥3 mice collected at 8 weeks of age for each experiment and is depicted by individual dots in each graph. Graphs are presented as mean +/− SEM and statistical significance is determined by an unpaired Student’s t-test. *p<0.05, **p<0.01, ***p<0.001 ns=not significant.

### Dek promotes expression of genes associated with luminal alveolar cells

Dek-OE correlated with a cellular ontology of “mammary gland luminal progenitor cells” and several upregulated genes are linked to milk protein production. To validate these transcriptomic profiles translated to increased protein expression, we performed western blot and immunohistochemical staining for luminal alveolar markers and milk proteins on whole mammary glands from 8-week-old virgin females. As predicted, Dek-OE mammary glands produced higher levels of milk proteins like Csn2 (β-casein; Fig 4A-B) and Muc1 (Mucin1; Fig 4C-D), even in virgin glands, and higher expression of luminal alveolar cell markers such as cytokeratins-7 and -19 and EpCAM (Fig 4E-F). However, markers of the luminal hormone receptor positive cell population, estrogen receptor alpha (ERα) and HER2, were unchanged in Dek-OE glands compared to controls (data not shown). This indicated that Dek function is limited in this population and that Dek expression is likely downstream of, or unrelated to, these luminal population molecular biomarkers that are often associated with breast cancers. Furthermore, whole mount Dek-OE mammary glands from 12.5 dpc pregnant mice demonstrated hyperplasia compared to controls (Fig 4G-H). The expanded cell population was morphologically similar to luminal alveolar cells, which expand during pregnancy. Since cellular expansion of the mammary gland is hormonally driven, and since *DEK* was previously published to be an ERα target gene in human cells, we determined whether endogenous Dek expression in the mammary gland was dependent on ovarian hormones. Immunohistochemistry staining revealed that Dek expression in the mammary epithelium was reduced by nearly 50% within one week of ovariectomy compared to intact controls (Fig 4I-J). This demonstrated that ovarian hormones, like progesterone and estrogen, partially promote Dek expression in mammary epithelium. We also investigated Dek expression across the developmental lifespan of the mouse mammary gland and show that Dek is highest during pregnancy and minimally expressed during lactation and involution (Fig 4K).

**Fig 4:**
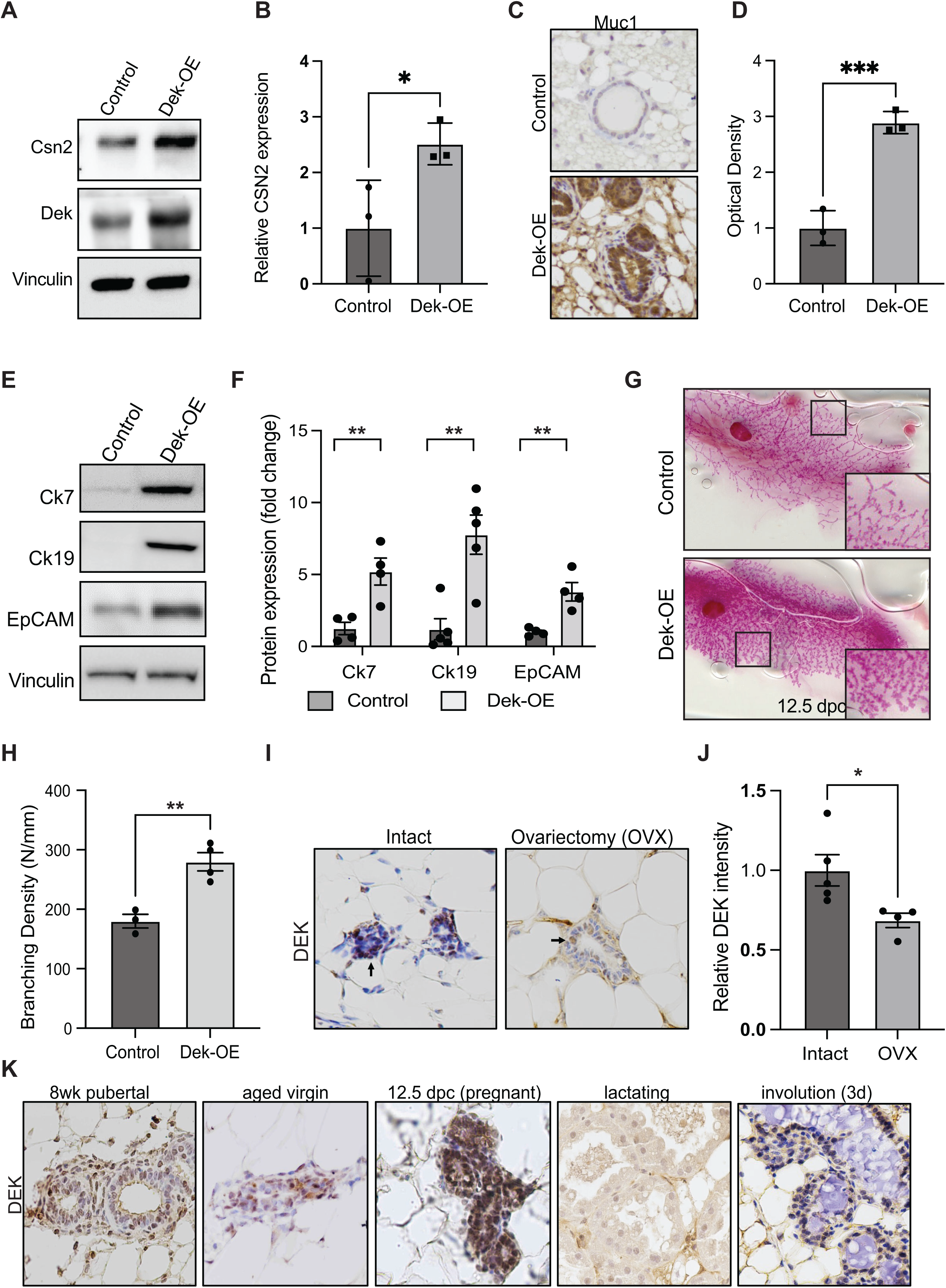
Dek overexpression expands the luminal-alveolar cellular compartment of the mammary epithelium. Dek over-expression correlates with expression of luminal alveolar markers and epithelial expression during pregnancy. (A) Dek-OE mammary glands express more Csn2 (β-casein) than controls as determined by western blotting of whole gland tissue. (B) Quantification of western blot images from (A) using densitometry. (C) Dek-OE mammary epithelium expresses significantly more Muc1 (Mucin1) compared to +dox controls as detected by immunohistochemical staining. (D) Quantification of staining intensity for Muc1 immunohistochemistry shown in (C). (E) Luminal alveolar markers cytokeratins -7 and -19 and EpCAM are highly expressed in whole mammary glands from Dek-OE mice compared to +dox controls, as detected by western blotting. (F) Densitometry quantification of western blots from (E). (G) Whole mount images reveal that Dek-OE results in increased expansion of mammary epithelium during pregnancy at 12.5 dpc. Specifically, the Dek-OE expansion in is the lobuloalveolar compartment. (H) Quantification of whole mounts from (G) using Sholl analysis via Image J. (I) Dek expression, detected by immunohistochemistry, in mammary epithelial cells is decreased one week after ovariectomy (OVX) compared to intact (non-OVX) female mice. (J) Quantification of staining intensity from (I) using Image J color deconvolution to detect DEK staining in epithelial cells. (K) Immunohistochemical staining shows that Dek expression levels are highest during pregnancy (12.5 dpc), moderate in epithelium from pubertal and aged virgin glands, and is nearly absent during lactation and involution. Graphs are presented as mean +/− SEM and statistical significance is determined by an unpaired Student’s t-test. *p<0.05, **p<0.01, ***p<0.001 ns=not significant.

We next analyzed *Dek* expression in a single cell murine mammary cell gene atlas, which was created by Saeki *et al.* by combining four single cell RNA sequencing datasets.^[35]^ *Dek* mRNA levels were highest in mammary stem cells (MaSC), luminal alveolar progenitors (LA-pro), and luminal hormone receptor positive (LH-pro) cells (Fig 5A). A subset of basal cells also expressed *Dek* while minimal expression was observed in mature, differentiated luminal cells. Importantly, these data support our bulk RNA-Seq data from Dek-OE mammary glands in that both datasets link *Dek* with luminal alveolar and stem/progenitor cell markers (see Fig 2). Gene set enrichment analysis (GSEA) of the genes most strongly correlated with *Dek* expression in the LA-pro cluster identified cell cycle (“E2F targets”), chromatin remodeling, and epigenetic regulation as the most enriched gene sets associated with *Dek* levels (Fig 5B, Fig S4-5). The 25 genes that were most strongly correlated with *Dek* are shown in Fig 5C. Cell clusters expressing the correlated epigenetic factor and histone methyltransferase, *Ezh2*, the proliferation marker *Pcna*, and the *E2F1* transcription factor that regulates the G1/S cell cycle transition, also overlapped with those expressing high levels of *Dek* (Fig 5D). Importantly, we noticed that two components of the PRC2 complex, *Ezh2* and *Rbpp4*, were strongly correlated with *Dek* expression and were highly expressed in the same three cell clusters as *Dek*: MaSC, LH-pro, and LA-pro cells (Fig 5E). Another PRC2 complex member, *Suz12* (correlation coefficient = 0.24), was also correlated with *Dek*, although it was not in the list of top 25 most correlated genes. The PRC2 complex utilizes the histone methyltransferase activity of Ezh2 to trimethylate histone H3 on lysine 27 (H3K27me3). Since *Dek* is known to function as a histone H3 chaperone in other systems, we tested whether Dek is a co-factor regulating histone H3 epigenetic post-translational modifications.

**Fig 5:**
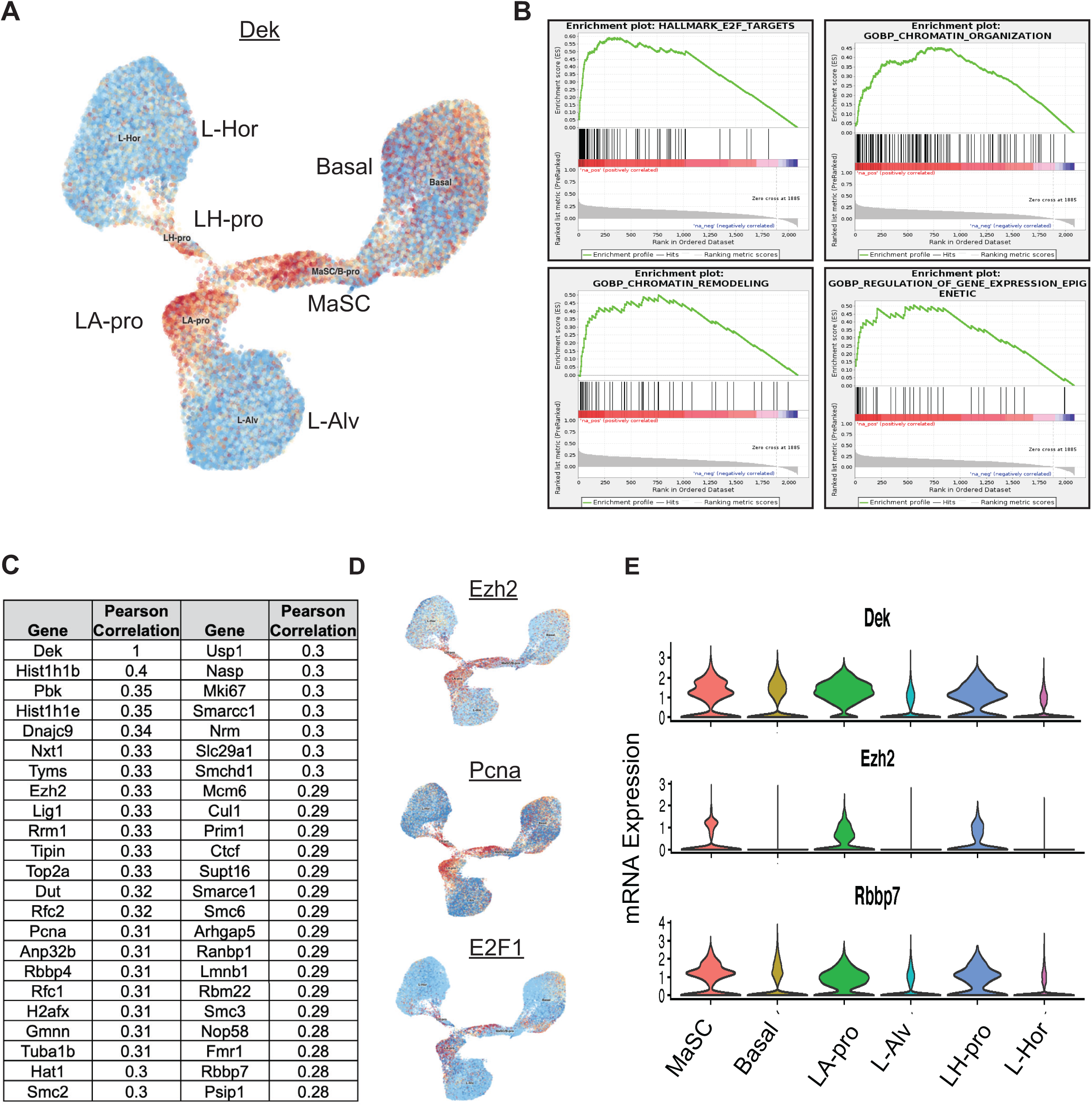
*Dek* is highly expressed in mammary stem cells and luminal progenitor cells and is co-expressed with genes associated with proliferation and chromatin remodeling. (A) UMAP plot depicting Dek expression (red) in single cell RNA-sequencing data from mouse mammary gland previously published by Saeki *et al.*^[35]^ The populations are mammary stem cells (MaSC), luminal alveolar progenitor cells (LA-pro), luminal hormone receptor positive progenitors (LH-pro), basal cells, mature luminal alveolar cells (L-Alv), and mature luminal hormone receptor positive cells (L-Hor). (B) Gene set enrichment analysis (GSEA) plots for genes co-expressed with Dek in the luminal alveolar progenitor cellular compartment. Plots depict E2F Targets, Chromatin Organization, Chromatin Remodeling, and Regulation of Gene Expresion-Epigenetic ontologies. (C) A list of the top genes co-expressed with Dek in the luminal alveolar progenitor cell population. (D) UMAP plots for gene expression of *Ezh2,* and proliferation-associated genes *Pcna* and *E2F1,* overlap with the expression of Dek in (A). All three genes are in the list shown in (C). (E) *Dek*, *Ezh2*, and *Rbbp7* mRNA expression levels in each mammary cell population are depicted as a violin plot. All three genes are most highly expressed in MaSC, LA-pro, and LH-pro populations.

### Dek increases H3K27 trimethylation in murine and human cells

DEK has previously been shown to repress histone H3 acetylation, which leaves these amino acid residues available for methylation.^[11, 14, 36]^ We quantified *Ezh2* mRNA expression in the bulk RNA-Seq samples. A 4.2-fold increase of *Ezh2* was observed in Dek-OE versus control mammary gland tissue (Fig 6A and 2E). A significant increase in H3K27me3 levels, relative to total histone H3, was observed by western blotting of mammary gland lysates (Fig 6B). We validated Ezh2 and H3K27me3 upregulation specifically in mouse mammary epithelial cells by immunohistochemistry (Fig 6C). To test if DEK-OE correlated with EZH2 and H3K27me3 in human cells, DEK was over-expressed or knocked-down in human MCF10A immortalized mammary epithelial cells. We observed a direct correlation, with H3K27me3 levels increased with DEK-OE and decreased in DEK-shRNA cells (Fig 6D). To probe a potential interaction between DEK and components of the PRC2 complex, a GFP-trap assay was performed with GFP-tagged DEK in HEK293 cells. GFP-DEK interacted with EZH2, RBBP4, and EED of the PRC2 complex as well as histone H3 (Fig 6E). CK2 was used as a positive control for a DEK interacting protein.^[37, 38]^ Notably, endogenous DEK was also pulled down in the GFP-trap assay, confirming that GFP-DEK can still multimerize with untagged DEK, as previously reported (Fig 6E).^[39]^ Additionally, endogenous DEK interacted with endogenous EZH2 and, potentially, SUZ12, in MCF10A mammary epithelial cells by immunoprecipitation (Fig 6F). Finally, to determine if DEK and EZH2 expression correlate in other models, we quantified their expression in breast cancer datasets. DEK and EZH2 protein (Fig 6G) and mRNA (Fig 6H) expression levels were, indeed, strongly positively correlated in human breast cancers. Interestingly, *DEK* and *EZH2* mRNA levels tended to be higher in estrogen receptor (ER) negative breast cancers (Fig 6H).

**Fig 6:**
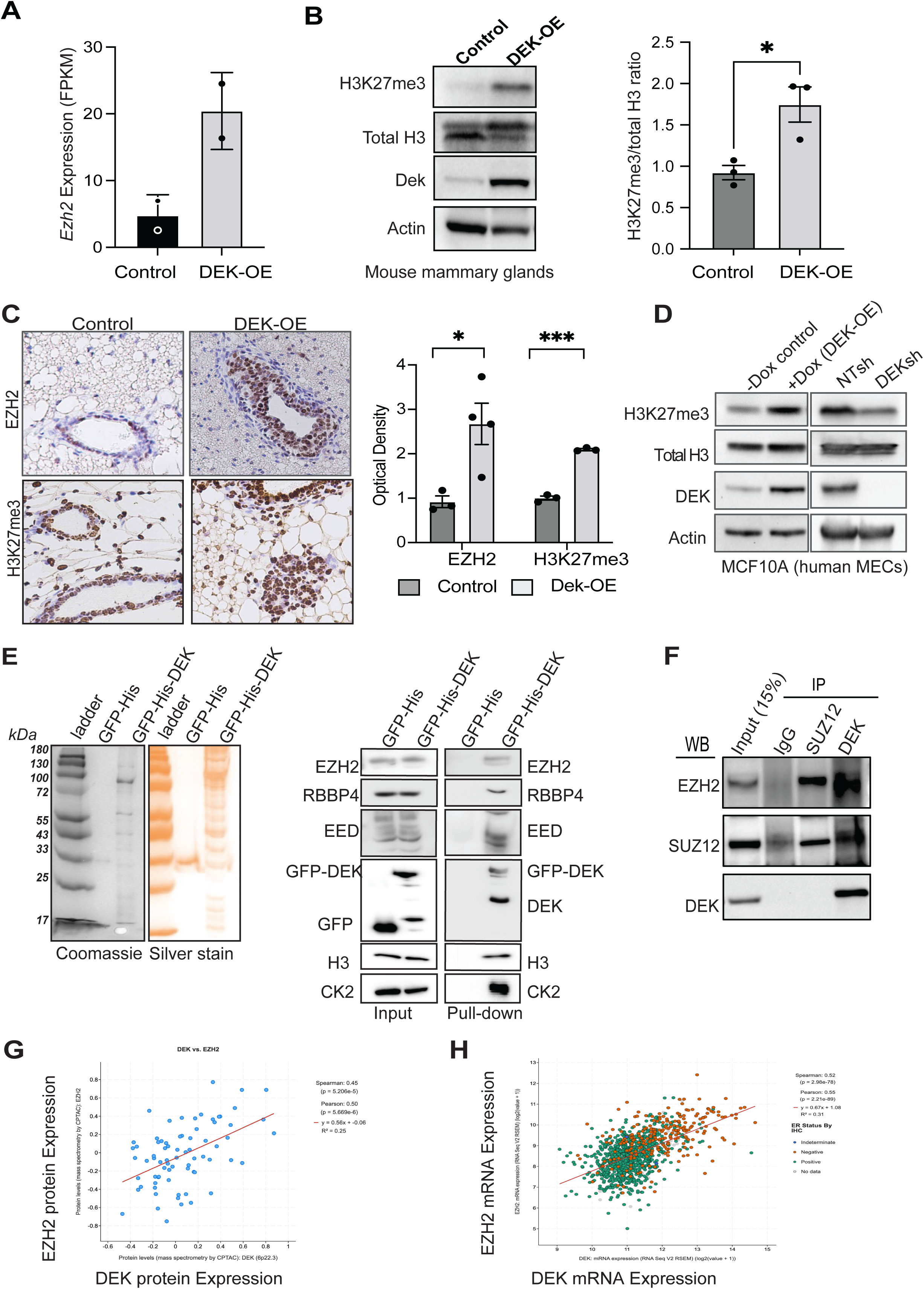
Dek interacts with the PRC2 complex to promote H3K27me3 epigenetic modifications in mouse and human cells. (A) *Ezh2* gene expression from bulk RNA-Seq of control versus Dek-OE mouse mammary glands, as described in figure 2. (B) Mammary glands from Dek-OE mice have increased levels of histone H3 trimethylation on residue lysine 27 (H3K27me3) compared to controls, as shown by western blot. Densitometry quantification is graphed to the right. N=3 mice per group. (C) H3K27me3 and EZH2 protein levels are increased in Dek-OE mammary epithelium from 8-week old female mice, compared to controls, as determined by immunohistochemistry. Staining intensity, quantified as optical density using Image J, is graphed to the right with N=3-4 mice/group. (D) H3K27me3 levels positively correlate with DEK expression in human MCF10A cells either over-expressing DEK (left) or with DEK knockdown with shRNA (right) as determined by western blot of whole cell lysates. (E) (left) Coomassie and silver stain of all proteins precipitated with GFP-DEK in a GFP-trap protein interaction assay compared to a GFP-His protein negative control performed in HEK293 cells. (right) Immunoblotting of GFP-trap nuclear lysates identify proteins that interact with GFP-DEK. Interacting proteins include key members of the PRC2 complex, EZH2, RBBP4, and EED, as well as endogenous DEK, histone H3, and the known DEK-interacting positive control of CK2. Input lysates, pre-incubation with GFP trap beads, are shown adjacent to the pull-down results. (F) DEK interacts with PRC2 complex members EZH2 and SUZ12 as determined by immunoprecipitation using whole cell lysates from MCF10A cells with dox-induced DEK overexpression. (G) EZH2 and DEK protein levels show a strong positive correlation in human primary invasive breast cancers. Pearson correlation=0.50, p=5.669e-6. (H) *EZH2* and *DEK* mRNA levels, detected by RNA-Seq, show a strong positive correlation in human primary breast cancers. Pearson correlation=0.55, p=2.21e-89. Estrogen receptor (ER) negative samples, as determined by immunohistochemistry, are shown as orange circles while ER positive samples are in green. Blue and gray dots are samples with indeterminate or no data, respectively. Data for (G) and (H) are from the TCGA Firehose Legacy dataset for breast invasive carcinoma, accessed using www.cbioportal.org, and graphed using a log scale. Trend lines are shown in red. Graphs are presented as mean +/− SEM and statistical significance is determined by an unpaired Student’s t-test. *p<0.05, **p<0.01, ***p<0.001 ns=not significant.

We next created a Dek-floxed mouse model and bred it to the CMV-Cre mouse to induce Dek deletion (Fig 7A). Dek protein loss was confirmed by western blot analysis of whole mammary tissue with a decrease noted for cytokeratin 7 expression, suggesting a decrease in the luminal cell population within the mammary gland (Fig 7B). Interestingly, at the time of weaning (21-28 days post-birth), litter sizes of Dek heterozygous and knockout dams were significantly smaller than litters of both wild-type (Dek^+/+^) and CMV-Cre-/Dek^fl/fl^ dams (Fig 7C). Survival analysis revealed that at least half of the pups born to Dek-deficient females died within 24 hours of birth (Fig 7D). Importantly, pup survival was not dependent on Dek genotype of the pups (data not shown) but was dependent on the genotype of the mothers. Pups born to Dek-deficient dams were dehydrated and lacked a milk spot, suggesting insufficient milk uptake (Fig 7E). Finally, we investigated if Dek loss impacted H3K27me3 levels, like Dek loss in human MCF10A cells (Fig 6D). Indeed, mammary glands from Dek knockout mice harbored significantly less H3K27me3 than controls by western blot analysis (Fig 7F-G) and immunohistochemistry (Fig 7I), that was accompanied by decreased levels of Ezh2 (Fig 7H). Combined, this indicates that mammary glands from Dek deficient mice are poorly functional and have impaired epigenetic remodeling, as evidence by decreased H3K27me3 levels. We conclude that Dek expression supports the proliferation of luminal alveolar progenitor cells in association with epigenetic reprogramming by stimulated EZH2 methyltransferase activity and histone H3 trimethylation at lysine 27 during mammary gland development.

**Fig 7:**
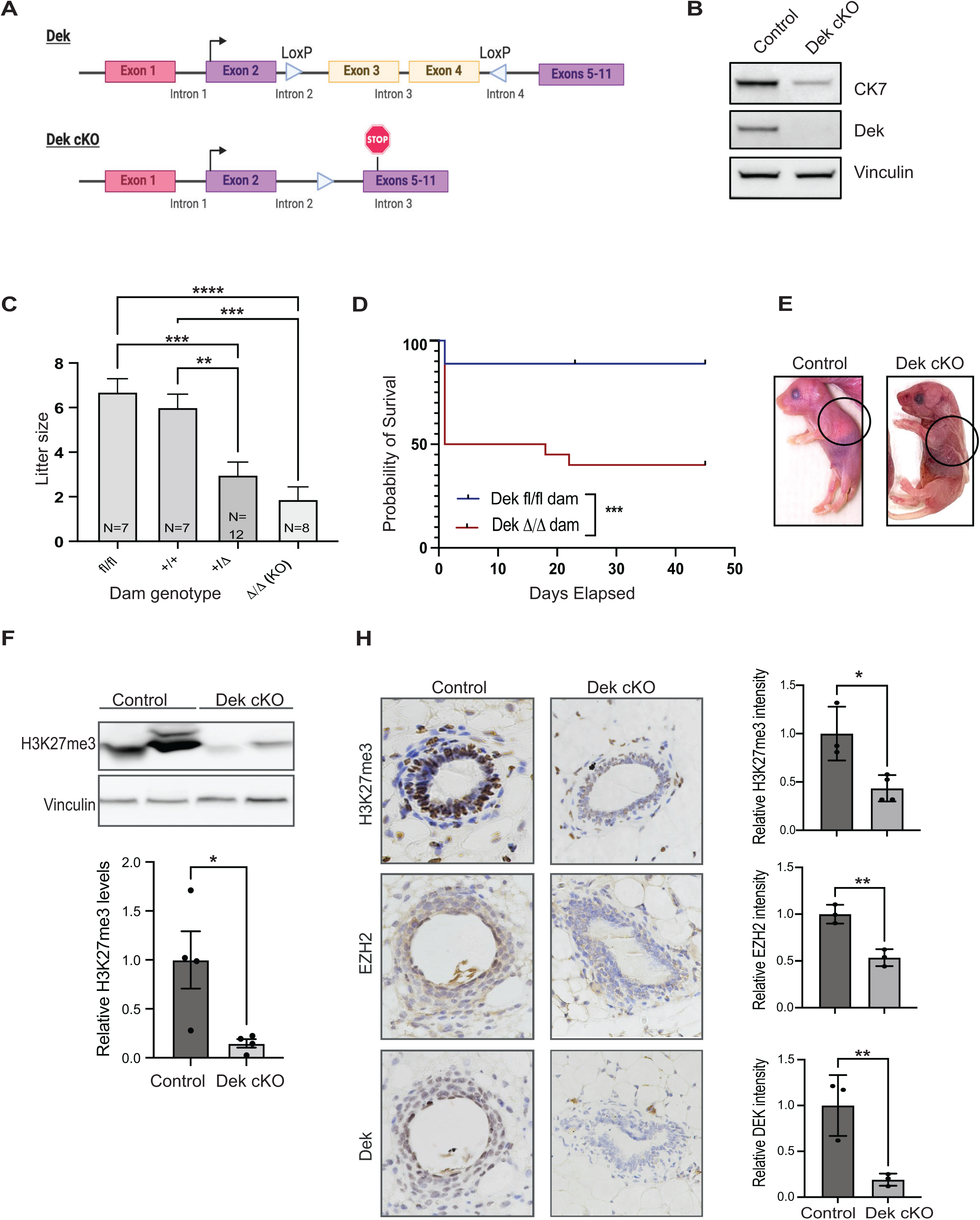
Mammary glands from Dek knockout mice are deficient in H3K27me3 and do not support neonatal survival. (A) Graphical representation of the floxed Dek allele in a novel conditional Dek knockout mouse model. LoxP sites flank exons 3 and 4 which, when removed by Cre recombinase, create a premature stop codon in exon 5. Exon 1 (pink) of Dek is non-coding. (B) Dek flox mice were bred to CMV-Cre mice to create a whole-body knockout. Whole cell lysates from mammary glands collected from 5-week-old female DEK knockout mice show loss of Dek expression and lower levels of luminal marker cytokeratin 7 (CK7). Vinculin is used as a loading control. (C) Dek knockout females have smaller litter sizes than Dek wild-type and CMV-Cre^−^/Dek^fl/fl^ controls. N values represent the number of separate dams in each Dam genotype group. N=7 for Dek^fl/fl^ and Dek^+/+^ controls, N=12 for Dek^+/Δ^ heterozygous dams, and N=8 for Dek^Δ/Δ^ knockout dams. (D) Kaplan-Meier survival curves reveal that half of pups born to Dek^Δ/Δ^ knockout dams (N=17 pups) die within 24 hours with a few more not surviving until weaning at 28 days. Minimal loss of pups are observed in litters born to Dek^fl/fl^ (N=20 pups). Data was collected from litters born to at least 3 separate dams per dam genotype. Statistical significance was determined by log-rank test (E) Photographs of pups born to Dek wild-type (WT, left) and knockout (KO, right) dams. Pups born to Dek WT dams had an observable milk spot within 24 hours. Deceased pups from Dek KO dams were dehydrated and did not have a milk spot. (F) Whole cell lysates from adult Dek^Δ/Δ^ mammary glands had significantly less H3K27me3 levels than mammary glands from adult Dek^fl/fl^ mice. (G) Densitometry quantification of western blot data from (F) as determined by Image J. N=4 per genotype. (H) Ezh2 expression and H3K27me3 levels are lower in the mammary epithelium of adult Dek^Δ/Δ^ mice compared to Dek proficient controls as determined by immunohistochemistry. Quantification of staining intensity are graphed on the right and presented as mean +/− SEM. Statistical significance is determined by an unpaired Student’s t-test. *p<0.05, **p<0.01, ***p<0.001 ****p<0.0001 ns=not significant.

## Discussion

Here, we are the first to describe a novel murine mammary gland model of temporal, tissue-specific Dek expression with transcriptomic profiling. We find that 2-fold upregulation of Dek promotes epithelial hyperplasia characterized by cell cycle deregulation. Many studies have shown DEK to be highly expressed in proliferating cells, as defined by co-expression between DEK and Ki67 or BrdU+ cells, but it was not clear if DEK expression could promote the cell cycle. Here we find adult Dek-OE mice display significantly more epithelium than their control counterparts, coinciding with an imbalance of cyclin A, CDK2, p21, and p27 that leans towards a pro-proliferative signaling mechanism. The cumulative life-time effect of discrepancies in cell cycle signaling resulted in hyperplastic mammary epithelium in Dek transgenic mice. Previously, the molecular mechanism(s) by which DEK promoted proliferation were not well defined. We note that cell cycle deregulation appears to begin from the early stages of cell cycle control, via down-regulation of cyclin/CDK inhibitors p21 and p27. This suggests that DEK expression may be a key component of exit from quiescence and entry into the G1 phase of the cell cycle and cell cycle progression. Importantly, DEGs in Dek-OE mammary glands were associated with both the cell cycle and “TP53 regulated transcription of cell cycle genes” (Fig 2F). This aligns with previous reports that DEK silencing by RNAi in glioblastomas and HeLa cells increases p53 upregulation and subsequent *CDKN1A*/p21 expression while exogenous DEK silences*TP53* (p53) expression.^[40–42]^ Other studies have also noted that DEK is associated with stem and progenitor cells exiting quiescence and induced cellular proliferation, including reports in murine muscle satellite cells, hematopoietic multipotent progenitors, human breast cancer stem cells, and *Artemia* crustaceans.^[25, 42–45]^ p53 is a key regulator of cell cycle control and quiescence, including maintaining quiescence in epithelial progenitor cells through the regulation of *CDKN1A/p21* expression.

Transcriptional control of *Cdkn1a/p*21 is positively regulated by p53 but both *Cdkn1a* and *Cdkn1b* are transcriptionally repressed via H3K27me3-mediated epigenetic silencing of their promoters.^[2, 46, 47]^ Histone H3 trimethylation on lysine 27 (H3K27me3) is accomplished through the methyltransferase activity of Ezh2 and the PRC2 complex. Here, we report for the first time that Dek is co-expressed with Ezh2 in murine and human tissues, promotes the creation of the H3K27me3 epigenetic mark, and physically interacts with PRC2 complex members EED, RBBP4, and EZH2.

Importantly, genetic mouse models revealed that regulators of H3K27me3 levels are essential for maintaining the luminal alveolar progenitor cell population and proper alveologenesis. Pregnancy results in a redistribution of H3K27me3 marks across the genome in luminal progenitor cells and increased EZH2 expression.^[48]^ Mice with mammary gland specific loss of EZH2 expression (MMTV-Cre/EZH2^f/f^) have a 14-fold *decrease* in gland repopulating mammary stem and progenitor cells, reduced alveolar cell development, and lactation failure.^[48]^ On the contrary, EZH2-OE in murine mammary glands results in hyperproliferation of luminal cells and delayed involution.^[49]^ These phenotypes closely resemble the ones we report here for DEK knockout and over-expression, respectively. Combined, it is evident that tightly regulated EZH2 expression, and H3K27me3 levels, are necessary for proper mammary alveologenesis during pregnancy, subsequent lactation, and involution of the gland at weaning. Our data indicates that Dek is an important component of this epigenetic regulation of mammary gland development and alveologenesis. Future studies will be required to determine how Dek may be involved in PRC2 complex recruitment to promoters and/or methyltransferase activity and how this intersects with p53 transcriptional activity. Our work suggests that Dek may prevent the expression of p53 target genes, like *Cdkn1a*, and others through H3K27me3 epigenetic silencing of their promoters. This has significant implications for pathogenesis within the mammary gland, since p53 is a potent tumor suppressor, frequently mutated in breast cancers, and DEK and EZH2 are both highly expressed in breast cancer with oncogenic activity, particularly in triple negative breast cancers.^[17, 18, 50–53]^ Although our Dek-OE transgenic mice did not form tumors through 15 months of follow-up, the hyperplasia phenotype suggests that Dek may support tumor progression, rather than initiation. The use of these genetic mouse models of mammary-specific Dek expression will be useful tools for understanding the role of Dek in breast cancer pathogenesis and mammary gland development *in vivo*.

Based on gene ontology analysis, Dek also up-regulated expression of multiple genes linked to the metabolism of RNA and proteins, with more specific cellular processes of nonsense-mediated decay, ribosomal RNA processing, and translation. While we did not investigate this directly in this manuscript, it is certainly worth investigating in the future. Interestingly, these cellular processes are supported by previous work that defined the DEK interactome in HeLa cells. DEK interacting proteins such as IMPDH2, DDX21, and numerous proteins of the large ribosomal subunit (RPL protein family) also implicate DEK in metabolism of RNA (nucleotide synthesis), ribosome RNA synthesis, and translation.^[37]^ Increased DEK expression has also been linked to AKT pathway activation, which regulates translation and exit from quiescence.^[12, 50, 54]^ Matrka *et al.* reported that DEK over-expression in epithelial cells reprograms metabolism to support macromolecule synthesis.^[55]^ With exit from quiescence and activation of the cell cycle, it is essential that the cell upregulates the production of nucleotides and amino acids to double their mass for eventual mitosis. More work is needed to understand how DEK expression impacts macromolecule synthesis and metabolic reprogramming.

Finally, we observed that Dek induced the expression of genes associated with lactation, including milk proteins, luminal alveolar cell markers, and stem/progenitor cell markers. Visually, whole mount analysis revealed hyperplasia that resembled the gland in early stages of pregnancy, which is when luminal alveolar progenitor cells are rapidly proliferating in preparation for lactation. Furthermore, a newly created conditional knockout mouse presents with decreased expression of luminal marker CK7 and poor pup survival prior to weaning. We also observed striking loss of H3K27me3 levels in Dek-deficient mammary glands. These findings are supported by sc-RNA-Seq data that revealed Dek expression is highest in mammary stem and progenitor cell populations in murine glands and that Dek and Ezh2 are co-expressed in these populations. Altogether, we are the first to report that Dek expression strongly supports mammary gland development, particularly as it relates to pregnancy, by promoting proliferation of stem and luminal progenitor cells. It is unclear how this impacts eventual lactation and the amount of milk produced to sustain pup viability in our mouse models, which is another limitation of this work. We have previously reported that Dek is an estrogen and progesterone target gene;^[16]^ therefore, it is possible that Dek facilitates epigenetic remodeling and transcriptional control of gene expression in response to elevated pregnancy hormones. It will be important to investigate this potential mechanism in future studies.

In summary, we report novel functions for Dek in promoting H3K27me3 epigenetic modifications through Ezh2 expression and interactions with the PRC2 complex. We also are the first to report that Dek supports mammary gland development through cell cycle control and, likely, RNA and protein synthesis. These findings have significant implications for understanding normal mammary gland development and disease pathogenesis, such as breast cancer.

## Materials and Methods

### Mice

Bi-L-Dek mice were kindly donated by Susanne Wells (Cincinnati Children’s Hospital Medical Center, Cincinnati, OH)^[29]^ and MMTV-tTA mice were kindly donated by Kay-Uwe Wagner (University of Nebraska Medical Center).^[31]^ Bi-transgenic mice were generated and maintained on an FVB/N background. DEK-OE mice were generated by continuously mating MMTV-tTA and Bi-L-DEK mice until homozygosity was achieved and copy number stabilized, as determined by quantitative PCR with genomic DNA. Handling of mice was performed with the approval of the Cincinnati Children’s Institutional Animal Care and Use Committee and approved under protocol 2020-0037 and 2023-0043. All mice were housed in specific pathogen-free housing with *ad libitum* access to food, with or without doxycycline, and water with a 12-h light/12-h dark cycle. DOX control animals were continuously fed DOX chow from maternal consumption during gestation through adulthood until the time of tissue collection.

The *Dek* conditional knockout allele was generated using CRISPR technology to introduce 5’ and 3’ loxP sites sequentially to flank exons 3 and 4. The sgRNAs were selected according to the on- and off-target scores from the CRISPR design web tool CRISPOR (http://crispor.tefor.net) ^[56]^. To insert the 5’ loxP site into intron 2, two sgRNAs (spacer sequences: GAGCTGTCAAGGTTACAGTG and GGGTTCCTGTAGGACATAG) were transcribed *in vitro* using the MEGAshorscript T7 kit (ThermoFisher) and purified by the MEGAclear Kit (ThermoFisher) and stored at −80C. To prepare the injection mix, sgRNAs (25ng/ul each) was mixed with Cas9 protein (IDT; 100ng/ul) in 0.1X nuclease-free TE buffer and incubated at 37C for 15 min to form the ribonucleoprotein complex (RNP). The donor oligo (Ultramer from IDT) with asymmetrical homologous arm design^[57]^ and the loxP sequence was added to the RNP at the final concentration of 100ng/ul. The zygotes from superovulated female mice on the C57BL6/N background (Taconic Biosciences) were injected with the mix via a piezo-driven cytoplasmic injection technique ^[58]^. Injected zygotes were subsequently transferred into the oviductal ampulla of pseudopregnant CD-1 females for birth. One pup with 5’ loxP in the litter was identified by PCR and Sanger sequencing and selected for breeding to homozygosity. To introduce the 3’ loxP site into intron 4, the zygotes from mice homozygous for 5’ loxP were collected and microinjected with Cas9 (100ng/nl), sgRNA (spacer sequence: TTCCTCTAGACCCAGTTAGG; 75 ng/ul) and a donor oligo (100ng/ul), followed by embryo transfer into pseudopregnant CD-1 females for birth. This resulted in two founder males carrying 5’ and 3’ loxP sites *in cis*, only one of which was able to sire progeny and establish the line. The line was back-crossed to C57Bl/6N for four generations to eliminate off-target genome editing and then re-bred to homozygosity, resulting in the conditional knockout Dek^fl/fl^ line that was maintained separately for future use. To test for phenotypes caused by Dek loss, Dek^fl/fl^ mice were bred to CMV-Cre mice on the C57Bl/6 background (Jackson Labs strain #006054). Once the deletion allele achieved germline transmission, the CMV-Cre transgene was eliminated from the colony and the strain was maintained as a constitutive knockout line via mating of heterozygous (Dek^+/Δ^) males and females.

Adult female C57Bl/6 mice underwent ovariectomy with isofluorane anesthesia. Pregnancy stage was determined by checking for plugs after placing a male and female in the same cage

### Genotyping

Tail clips were digested with DirectPCR Lysis Reagent (Viagen Biotech) containing 0.6mg/mL Proteinase K (Invitrogen) and protocol (need to read bottles/thermocycler). For PCR analysis one microliter of DNA was added to JumpStart Taq Ready mix from Invitrogen (Carlsbad, CA, product # P2893) using the manufacturer’s specifications.

Transgenes were detected with the following primers:

Bi-L-Dek transgene (DEK 3’): (F):GAAATGTCCGTTCGGTTGGCAGAAGC (R):CCAAAACCGTGATGGAATGGAACAACA. Luciferase: (F): AGTCGATGTACACGTTCGTCAC (R): TGACGCAGGCAGTTCTATGC tTA: (F): GCTGCTTAATGAGGTCGG (R): CTCTGCACCTTGGTGATC DEK (endogenous): (F) TCGAAATGCCATGTTAAAGAGCA (R): AAGGCTTTGGATGCATTAAGAAGT

DEK conditional knockout models were genotyped with the following primers: CMV-Cre: (F): GCGGTCTGGCAGTAAAAACTATC (R): GTGAAACAGCATTGCTGTCACTT

DEK (detects endogenous and floxed alleles): 5’loxP (F): AGTGAAATTACTGGTCTGTGAAG (R): CTGAGTGGAACAGCTCCTATAG 3’loxP (F): AGATGCTTCACCTTAGAGCTG (R): TCAGTTTGGAGCAAATTTCATTTCC DEK deletion allele: (5’loxP F): AGTGAAATTACTGGTCTGTGAAG (3’loxP R): TCAGTTTGGAGCAAATTTCATTTCC

### In vivo Imaging Systems (IVIS)

Mice were intraperitoneally injected with 15ng/g of luciferin and allowed to metabolize the luciferin for five minutes prior to sedation with inhaled isoflurane. Mice were imaged in the Perkin Elmer IVIS Spectrum CT, Waltham, Massachusetts, USA.

### Western Blotting

Mammary gland tissues flash frozen in liquid nitrogen and stores at −80C until ready to use. Tissues were homogenized using mortar and pestle with liquid nitrogen, transferred to 1.5ml tube and weighed, with 5μl of RIPA added for every 1mg of tissue. RIPA consisted of 1% Triton, 24mM sodium deoxycholate, 0.1% SDS, 0.16M NaCl, 10mM Tris pH 7.4, 5mM EDTA, 10mM Na_3_VO_4_, and 10mM NaF, supplemented with a protease inhibitor cocktail (Sigma Aldrich). Tissue was lysed on ice in a 4C cold room for 4 hours with vortexing every hour. Samples were sonicated 3x for 15 seconds with the amplitude set to 35. Tissue was incubated on ice an additional 20 minutes then cleared by centrifugation at 13,000rpm for 25 minute. Clearing by centrifugation and transfer to new tubes was performed three times to fully remove lipids. Protein quantifications were determined with a Bradford assay then 30μg of protein was separated on a 8-16% or 12% pre-cast gel (Bio-Rad) using primary antibodies diluted 1:500-1:1000 and secondary antibodies (1:2500; Cytiva). See Table S1 for list of antibodies used in this work. Membranes were exposed to enhanced chemiluminescence reagents (Thermo Scientific) and imaged using the BioRad Chemidoc (Hercules, CA, USA). Densitometry was performed using Image J.

### Mammary Gland Whole Mounts

Mice were sacrificed and the fourth inguinal mammary glands were excised, spread on a cover slip and prepared for whole mount fixation and staining as previously described.^[59]^ Whole mounts were imaged via Nikon Eclipse Ci upright microscope with Nikon Digital Sight camera and Nikon NIS-Elements software or digitally imaged with a flatbed scanner for Sholl analysis as previously described.^[60]^

### Immunohistochemistry

Tissues were fixed in 4% paraformaldehyde, embedded in paraffin, sectioned at 4 μm thickness, and fixed onto slides. Routine H&E-stained sections were analyzed for histopathology. Paraffin sections were deparaffinized in xylene and rehydrated for antigen retrieval in sodium citrate. Off-target background staining was reduced by pre-treatment with 0.3% H_2_O_2_. Sections were then treated with the Mouse on Mouse peroxidase immunostaining kit (Vector Labs, Burlingame, CA, USA) with primary antibodies. Sections were treated with 1:500 dilutions of biotinylated secondary antibody and stained with diaminobenzidine (DAB) and counterstained with hematoxylin then mounted with Permount (Fisher Scientific, Pittsburgh, PA, USA). Images were captured at the indicated magnifications with a Nikon Eclipse Ci upright microscope with Nikon Digital Sight camera and Nikon NIS-Elements software. Image J color deconvolution was utilized to measure the staining intensity only within mammary epithelial cells from at least 3 fields of view from at least 3 different mice per group.

### Cell culture

#### Primary Mammary Epithelial Cell Isolation and Culture

Mammary glands were dissected and epithelial cells were isolated via differential centrifugation as previously described.^[61]^ Primary mammary epithelial cells (MECs) were cultured in Epicult-B media with proliferation supplement (Stemcell Technologies, #05610) with 10ng/ml EGF, 10ng/ml bFGF, 4ug/ml Heparin, and penicillin/streptomycin. For organoid cultures, 8-chambered slides were coated in 100% Matrigel, then primary MECs were trypsinized and resuspended in 2.5% Matrigel in 50:50 mix of Epicult-B and 3D assay media at a density of 5000 cells per chamber as previously described.^[62]^ Palbociclib, or diluent (H_2_O), was added at a concentration of 1μM 72 hours after seeding cells in 3D culture, then organoids were imaged 96 hours later. Cells isolated from mice fed doxycycline chow were maintained in culture with 1μg/ml doxycycline.

#### Human Cell Lines

MCF10A human immortalized mammary epithelial cells were acquired from ATCC and cultured as previously described in DMEM:F12 medium with 5% horse serum, EGF, hydrocortisone, cholera toxin, insulin, and penicillin-streptomycin.^[62]^ DEK over-expressing cells were created by transduction with a modified doxycycline-inducible pTRIPZ lentiviral vector into which full-length, untagged, DEK cDNA had been cloned using the SnaBI and EcoRI restriction sites. DEK knockdown cells were created by lentiviral transduction of pLKO.1-DEK shRNA (Sigma MISSION shRNA #TRCN0000013104). Selection of transduced cells was performed with 2μg/ml puromycin and DEK expression induced with 1μg/ml doxycycline. HEK293 FLp-In T-Rex cells were cultured in DMEM with 10% FBS and 1% Pen–Strep 37°C, 95% humidity and 5% CO_2_.

### LAP-Tag immunoprecipitation

This assay was performed as previously described and with manufacturer’s instructions (Flp-Ub T-Rex Core kit, Invitrogen, #K6500-01).^[37, 63]^ Briefly, HEK293 cells were transfected with Lipofectamin 2000 and 2 µg of pcDNA5/FRT/TO DEK-His-GFP, or pcDNA5/FRT/TO His-GFP control, and 18 µg of pOG44. After analysis of expression (data not shown) HEK293 cells containing the inducible LAP-tag DEK fusions were transferred from six 10 cm plates into a spinner flask containing 250 mL DMEM/10% FBS in the presence of Pen/Strep then expanded to a total of 1L (roughly 10^6^ cells/mL). 24 hours prior to immunoprecipitation, expression of the LAP-tag fusion was induced with 50 ng/mL tetracycline. Nuclear extracts were prepared as described^[37]^ while 100µL GFP-trap beads was washed three times with 1 mL wash buffer (10mM Tris/HCl pH 7.5, 150mM NaCl, 0.5mM EDTA). The nuclear extract was added to the washed GFP-trap beads and incubated for 1 hour on a rolling platform at 4°C. The beads were washed three times with 1 mL wash buffer and eluted three times with 100µL 0.2M glycine pH 2.5 and neutralized in 1M Tris pH 10.4. The proteins were precipitated according to the Wessel/Flügge protocol^[64]^ and resuspended in 100μl 2% SDS. 2-4μl of eluates were analyzed by SDS-PAGE electrophoresis, followed by either Coomassie or silver staining or immunoblotting with the indicated antibodies. All LAP IP buffers were freshly supplemented with 5mM DTT, 0.1mM PMSF, and 1× Complete Protease inhibitor mix. For immunoblotting, the following antibodies were used: DEK (A0315, Abclonal, 1:1000), GFP (AF1483, Beyotime, 1:1000), CK2 (A19683, Abclonal, 1:1000), histone H3 (AF0009, Beyotime, 1:1000), RBBP4 (A3645, Abclonal, 1:1000), EED (A5371, Abclonal, 1:1000), EZH2 (A16846, Abclonal, 1:1000).

### Co-Immunoprecipitation

DEK expression in MCF10A pTRIPZ-DEK cells was induced by treating cells with 1μg/ml doxycycline for 48 hours prior to collection in IP lysis buffer (25mM NaCl, 25mM Tris-HCl, 1mM EDTA, 1% Nonidet P-40, and 5% glycerol) supplemented with phosphatase and protease inhibitors as in the western blotting section. Lysates were cleared with protein A/G beads and 1 mg of protein was incubated with 2 μg primary antibody (or 1:50 dilution if concentration was unknown) overnight with rotation at 4°C. Antibodies for DEK (Abcam #ab245429) or SUZ12 (D39F6) (Cell Signaling Technology, #3737) were used for immunoprecipitation. Antibody-protein complexes were precipitated with protein A/G agarose beads for 3 hours, washed, and removed from beads by boiling in 2x SDS-PAGE sample buffer with 6% β-mercaptoethanol then run on a 4-15% gradient gel for immunoblotting as described above. DEK (BD Biosciences #610948, 1:1000), SUZ12 (Cell Signaling Technology, #3737, 1:1000), and EZH2 (Proteintech, #21800-1-AP, 1:2000) primary antibodies were used to probe for co-immunoprecipitation.

### RNA sequencing

Mammary glands were collected from nulliparous aged female mice, three DEK-OE and three DEK-OE on dox chow since birth. RNA was isolated from the mammary gland with Trizol and approximately 80–120ng of RNA was amplified to generate cDNA. The initial amplification step for all samples was performed with the NuGEN Ovation RNA-Seq System v2 (NuGEN,San Carlos CA, USA). The concentrations were measured using the Qubit dsDNA BR assay. cDNA size was determined by using a DNA1000 Chip. Libraries were then created for both samples. Specifically, the Nextera XT DNA Sample Preparation Kit (Illumina, SanDiego, CA, USA), was used to create DNA library templates from the double-stranded cDNA. Concentrations were measured using the Qubit dsDNA HS assay. Then 1ng of cDNA was suspended in Tagment DNA buffer. Tagmentation (fragmentation and tagging with the adaptors) was performed with the Nextera enzyme (AmpliconTagment Mix, Illumina) by incubation at 55°C for 10 min. NT buffer was then added to neutralize the samples. Libraries were prepared by PCR with the Nextera PCR Master Mix and 2 Nextera Indexes(N7XX and N5XX) according to the following program: 1 cycle of 72°C for 3 min; 1 cycle of 98°C for 30 sec; 12 cycles of 95°C for 10 sec, 55°C for 30 sec, and 72°C for 1 min; and 1 cycle of 72C for 5 min. Purified cDNA was captured on an Illumina flow cell for cluster generation. The size of the libraries for each sample was measured using the Agilent HS DNA chip (Agilent Genomics, Santa Clara CA, USA). Libraries were sequenced on the IlluminaHiSeq2500 following the manufacturer’s protocol, with 75-bp paired-end sequencing and a coverage of 30M reads.

Quantification of mRNA expression levels was based on the TopHat/Cufflinks pipeline of the CCHMC DNA sequencing and Genotyping Core. Paired-end reads were aligned to mouse genome build mm10 with STAR version 2.6.1. Principal component analysis (PCA) highlighted potential batch effects resulting in the removal of one control sample and its paired DEK-OE sample for a final sample size of two per group. Transcripts per million (TPM) were generated using the pseudo aligner Kallisto and processed through the R package NOISeq version 2.42.0 for batch correction. AltAnalyze version 2.1.4.4 (EnsMart72 database) was used for differential expression analysis on the batch-corrected data matrix (Table S2). The AltAnalyze supplied empirical Bayes moderated *t*-test was performed followed by Benjamini–Hochberg adjustment for false discovery (FDR). Pathway enrichment was performed on genes that met a final threshold cutoff of a raw *p*-value ≤ 0.05 and fold change ≥ 1.5 utilizing the publicly available web-based tool, g:Profiler (https://biit.cs.ut.ee/gprofiler/gost accessed on 5 April 2023). g:Profiler results were visualized in node maps created in Cytoscape version 3.9.1. The volcano plot was created with the Enhanced Volcano package in R version 4.2.1 (https://github.com/kevinblighe/EnhancedVolcano) using all expressed genes. Genes with a raw p-value ≤ 0.05 upregulated ≥ linear 1.5-fold are indicated in red and genes significantly downregulated ≤ linear −1.5-fold genes are indicated in blue. A subset of highly regulated or mammary gland-related genes are highlighted in boxes. The raw and processed RNA-sequencing data are available upon request.

### scRNA-Seq analysis

Data previously published by Saeki et al. was obtained and the log normalized gene expression values from the six cell clusters was used for analysis.^[35]^ To identify genes co-expressed with Dek, only genes that were expressed in at least 50% of the cells in each cluster were included. Cells expressing Dek and the selected genes underwent Pearson correlation analysis. Cells that did not express Dek or the selected gene for comparison were removed from analysis. The Pearson correlation coefficient was used as a rank for each gene in gene set enrichment analysis (GSEA). GO and Hallmark gene sets were used for GSEA.

### Statistics

Error bars depict standard errors of data collected from at least three mice. Significance was set at p<0.05 (*p<0.05, **p<0.01). Log-rank test, Student’s t-test or one-way ANOVA was utilized to test for significance using GraphPad Prism.

## Acknowledgements

We thank Veterinary Services, Jeff Bailey and the Comprehensive Mouse and Cancer Core, the Gene Expression Core, and the DNA Sequencing Core facilities at Cincinnati Children’s Hospital Medical Center. Thank you to Jonathan Cheek and Jordan Harris for technical assistance. Funding for this work was provided by the Cancer Survivorship pilot grant from the University of Cincinnati Cancer Center (LMPV), a Trustee Award from Cincinnati Children’s Hospital (LMPV), National Cancer Institute awards T32CA117846-13 (MJ), and R37CA218072 (LMPV).

## Author Contributions

Data acquisition, data analysis, project development, and written drafts were completed by MJ and AL. Data acquisition and analysis was also completed by LGP, AL, TL, KW, and MSS. FK performed the LAP-tag immunoprecipitation and provided feedback on written drafts. Bulk and scRNA-Seq analysis was completed by TL, NS, and AP. Project development, conceptual design, data interpretation, funding, and written drafts were completed by LMPV with assistance from SIW. Funding was provided by MJ and LMPV.

---

**Fig S1:**
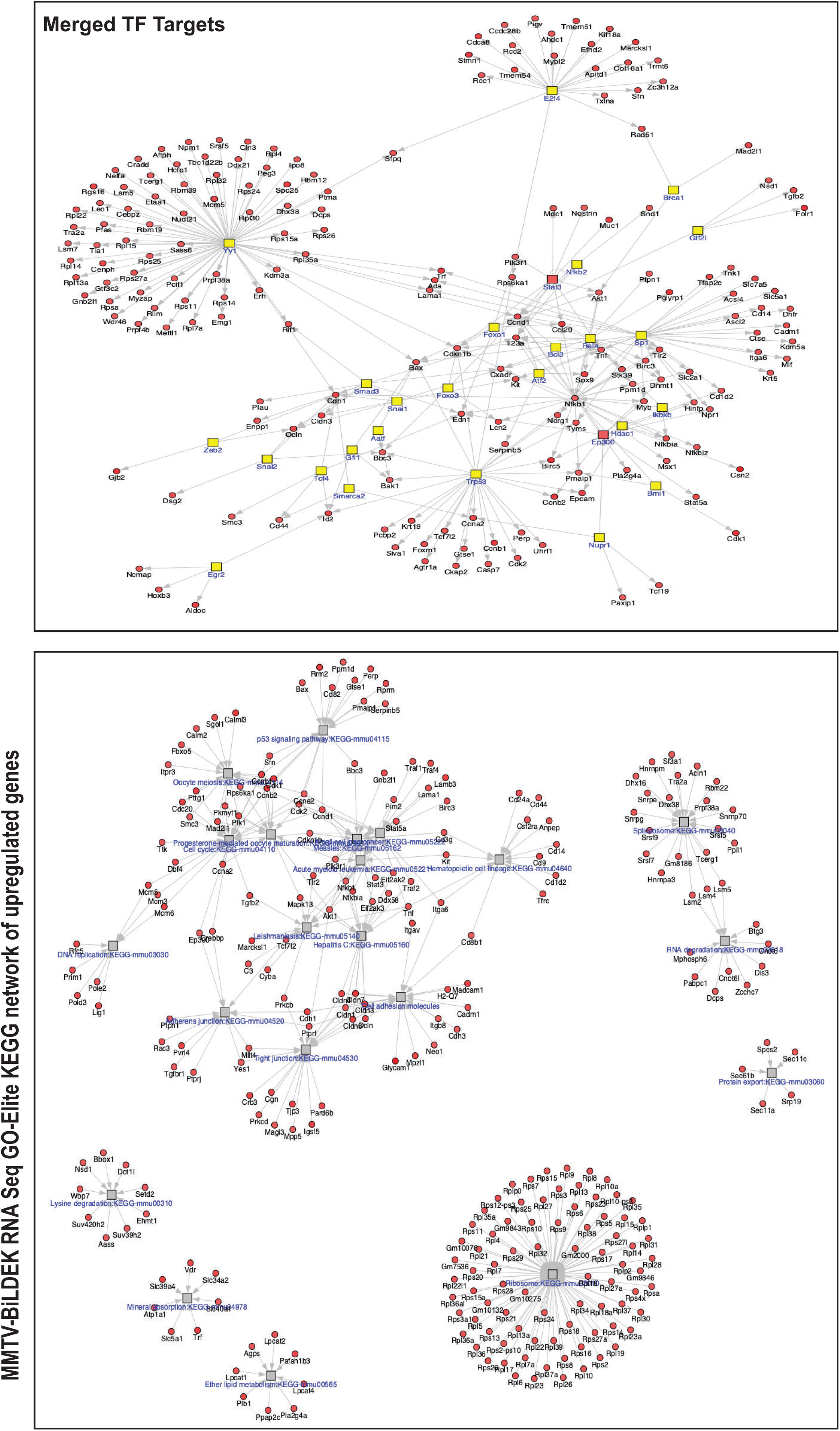
Analysis of up-regulated genes in Bi-L-Dek transgenic mammary glands compared to controls. (A) GO-Elite was used to identify transcription factors Yy1, Nfkb2, p53, and E2F as potentially responsible for the expression of Dek-OE-induced differentially expressed genes. (B) GO-Elite was used to assess KEGG pathways associated with DEGs caused by Dek over-expression. For upregulated genes, this identified p53 signaling, cell cycle, tight junctions and adherens junctions, ribosomes, and progesterone signaling.

**Fig S2:**
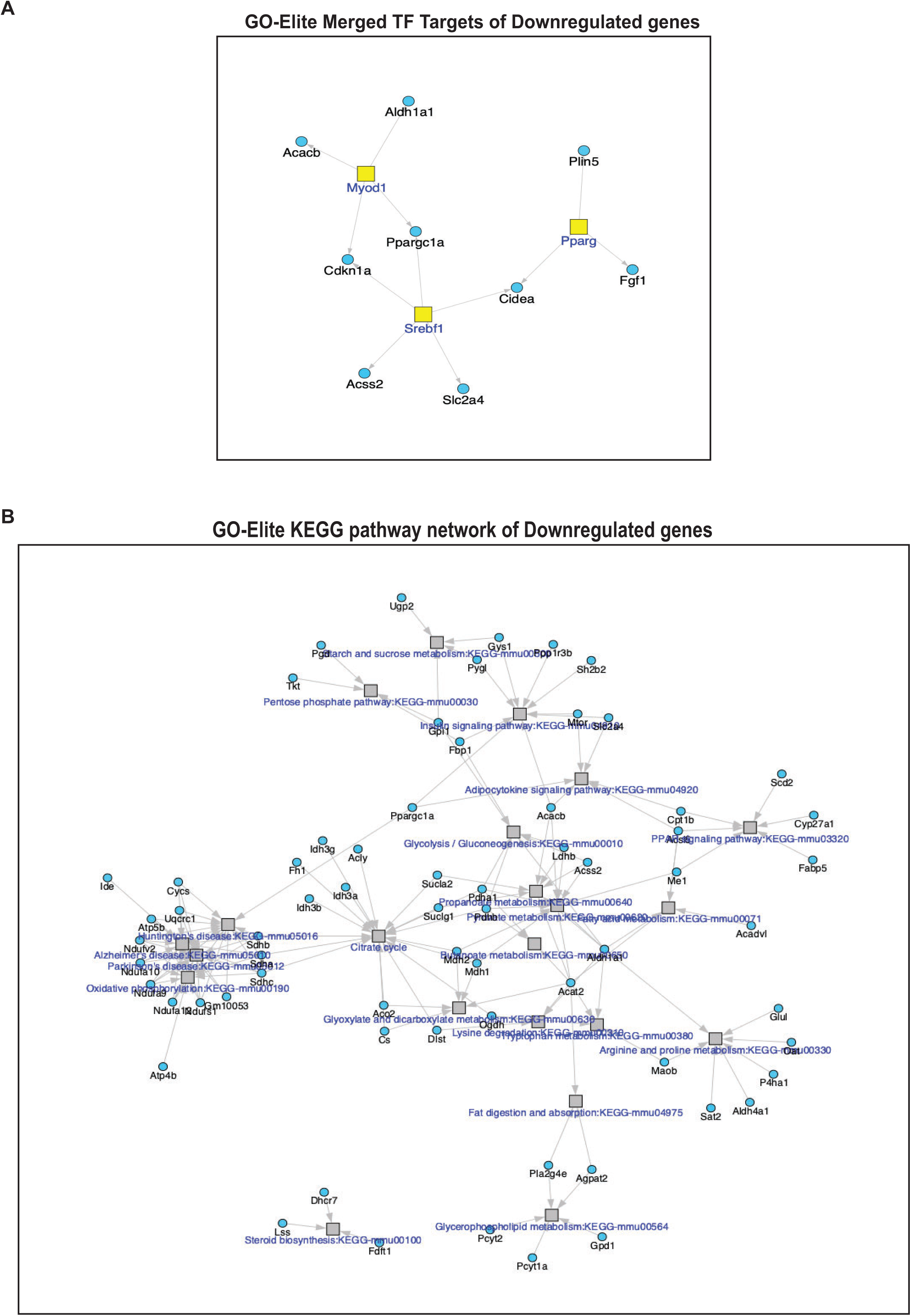
Analysis of down-regulated genes in Bi-L-Dek transgenic mammary glands compared to controls. (A) GO-Elite was used to identify transcription factors associated with the expression of down-regulated genes, which included Pparg, Myod1, and Srebf1 (B) For down-regulated genes, KEGG pathways were identified that related to adipocytokine signaling and a small number of genes related to various metabolic pathways like glycolysis, citrate cycle, and pentose phosphate pathways.

**Fig S3:**
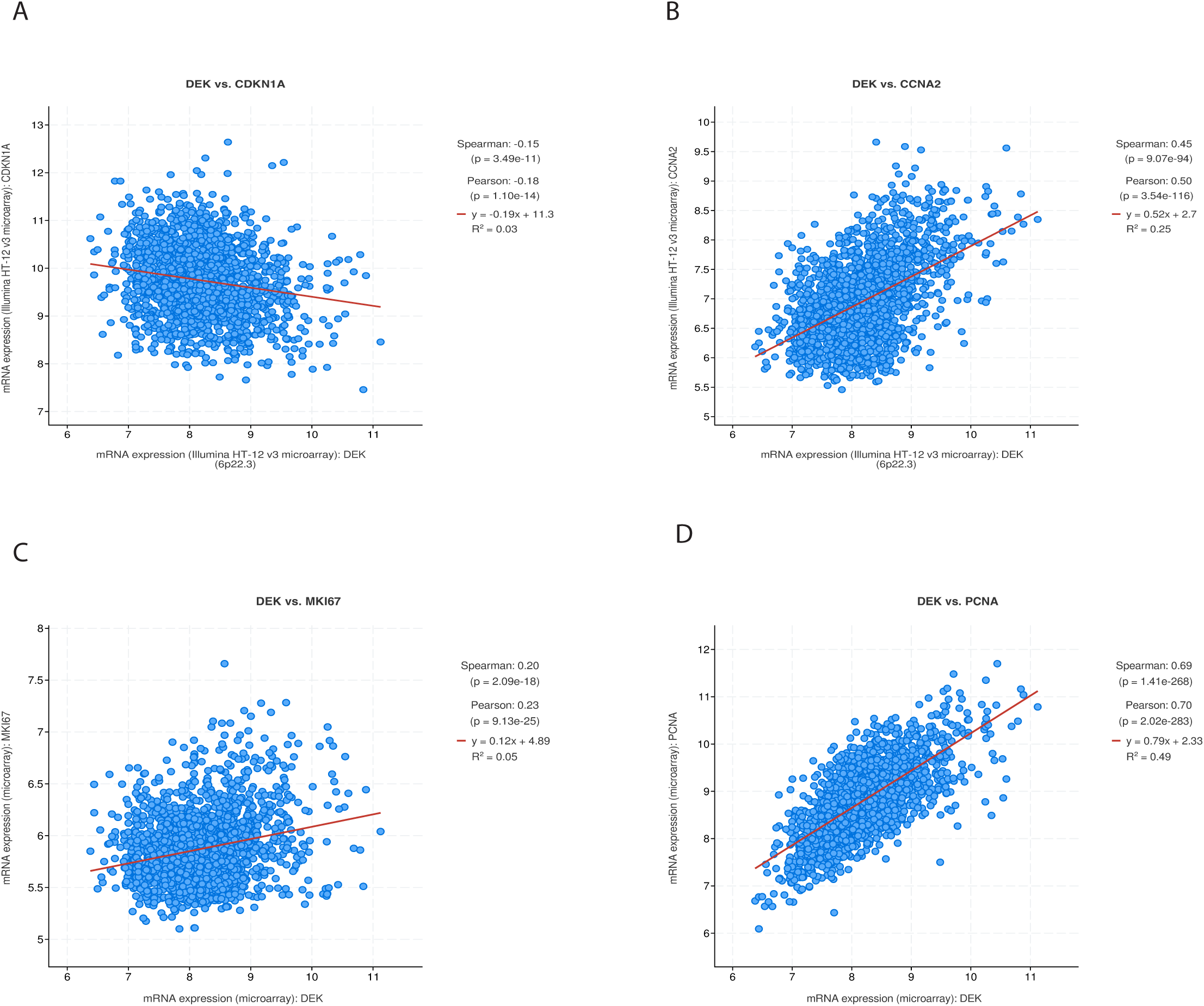
*DEK* correlates with cell cycle associated genes in human breast cancers. Genes co-expressed with *DEK* in human breast cancers were identified in the METABRIC dataset. *CDKN1A* (p21) was negatively correlated while *CCNA2* (Cyclin A2), *MKi67* (Ki67), and *PCNA* were positively correlated. Data accessed using CBioPortal (cbioportal.org).

**Fig S4:**
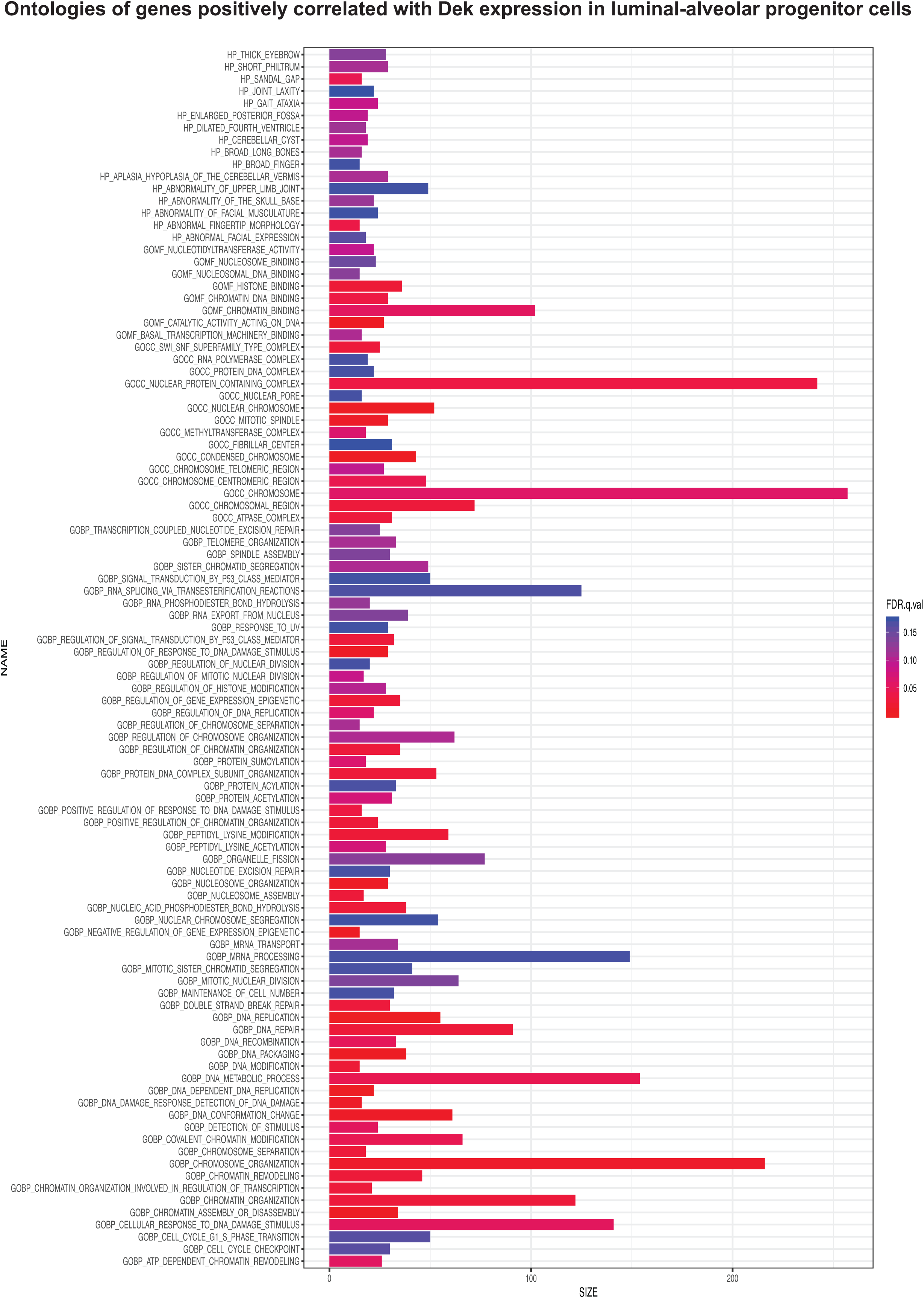
Ontologies of genes positively correlated with Dek expression in luminal alveolar progenitor cells of the mouse mammary gland. Gene set enrichment analysis (GSEA) of the genes most strongly positively correlated with *Dek* expression in the LA-pro cluster identified cell cycle (“E2F targets”), chromatin remodeling, and epigenetic regulation.

**Fig S5:**
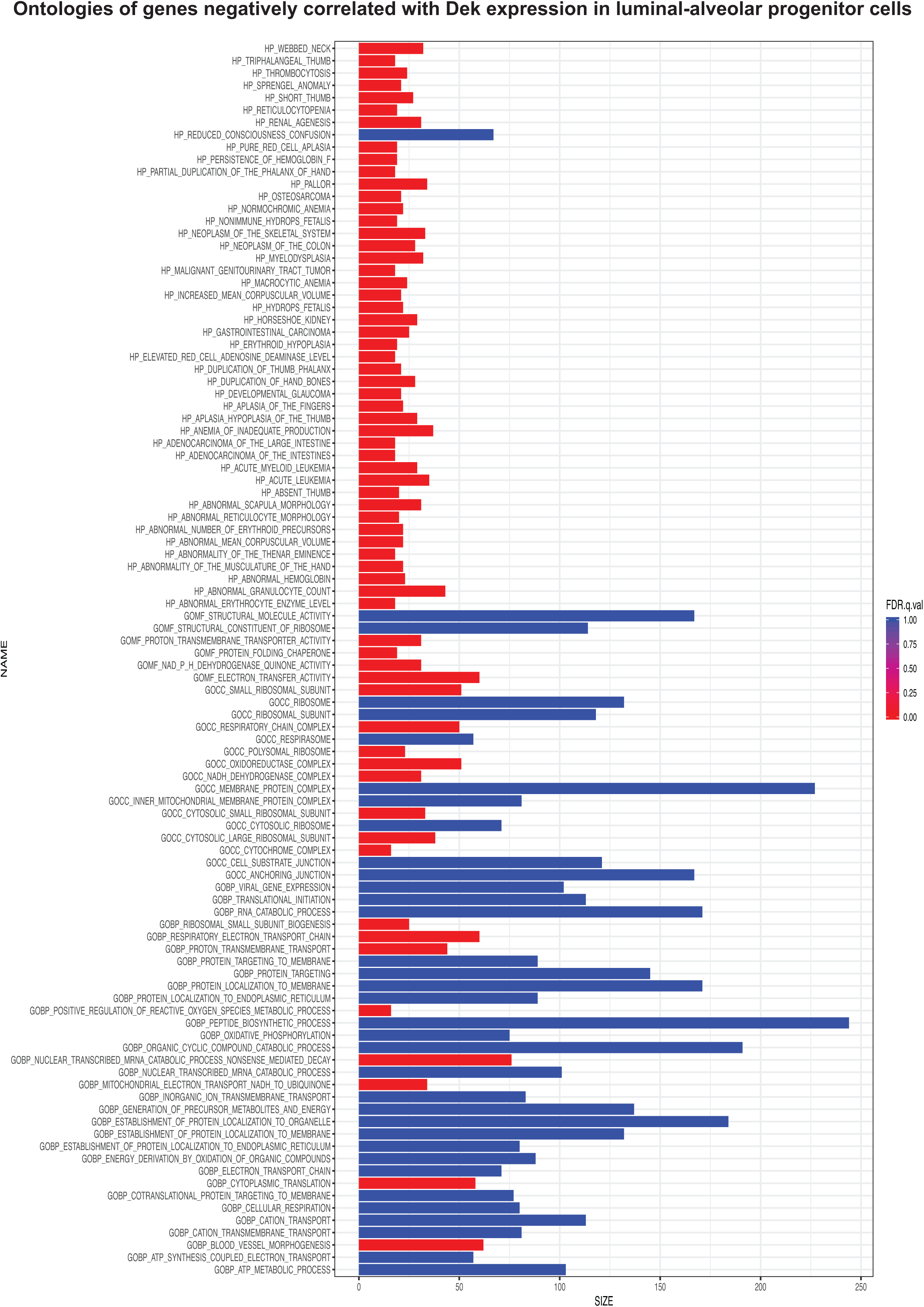
Ontologies of genes negatively correlated with Dek expression in luminal alveolar progenitor cells of the mouse mammary gland. Gene set enrichment analysis (GSEA) of the genes most strongly negatively correlated with *Dek* expression in the LA-pro cluster identified electron transport chain, protein folding, and other molecular functions.

**Table S1:** List of antibodies and dilutions used in this manuscript.

| Antibody Name | Manufacturer | Catalog # | Dilution (IHC/WB) |
| --- | --- | --- | --- |
| DEK | BD Transduction | 610948 | 1:100/1:1000 |
| CDK4 | Cell Signaling | 12790T | 1:100/1:1000 |
| CDK6 | Cell Signaling | 3136T | 1:100/1:1000 |
| MUC1 | invitrogen | PIPA595191 | 1:350/1:1000 |
| EZH2 | Cell Signaling | 5246 | 1:200/1:1000 |
| p21 | Abcam | ab188224 | 1:100/1:1000 |
| p27 | Cell Signaling | 3686 | 1:100/1:1000 |
| CSN2 | Fisher Scientific | 501556761 | 1:100/1:1000 |
| H3k27me3 | Cell Signaling | 9733 | 1:500/1:1000 |
| CDK2 | Cell Signaling | 2546 | 1:100/1:1000 |
| Vinculin | Cell Signaling | 13901S | 1:100/1:1000 |
| CK19 | Fisher Scientific | PIMA531977 | 1:100/1:1000 |
| CK7 | Abcam | ab9021 | 1:100/1:1000 |
| EpCAM | Bioss Antibodies | bs-1513R-FR | 1:100/1:1000 |

